# Anticipating future SARS-CoV-2 variants of concern through *ab initio* quantum mechanical modeling

**DOI:** 10.1101/2021.11.25.470044

**Authors:** Marco Zaccaria, Luigi Genovese, Michael Farzan, William Dawson, Takahito Nakajima, Welkin Johnson, Babak Momeni

## Abstract

Evolved SARS-CoV-2 variants are currently challenging the efficacy of first-generation vaccines, largely through the emergence of spike protein mutants. Among these variants, Delta is presently the most concerning. We employ an *ab initio* quantum mechanical model based on Density Functional Theory to characterize the spike protein Receptor Binding Domain (RBD) interaction with host cells and gain mechanistic insight into SARS-CoV-2 evolution. The approach is illustrated via a detailed investigation of the role of the E484K RBD mutation, a signature mutation of the Beta and Gamma variants. The simulation is employed to: predict the depleting effect of the E484K mutation on binding the RBD with select antibodies; identify residue E484 as a weak link in the original interaction with the human receptor hACE2; and describe SARS-CoV-2 Wuhan strand binding to the bat *Rhinolophus macrotis* ACE2 as more optimized than the human counterpart. Finally, we predict the hACE2 binding efficacy of a hypothetical E484K mutation added to the Delta variant RBD, identifying a potential future variant of concern. Results can be generalized to other mutations, and provide useful information to complement existing experimental datasets of the interaction between randomly generated libraries of hACE2 and viral spike mutants. We argue that *ab initio* modeling is at the point of being aptly employed to inform and predict events pertinent to viral and general evolution.

## 1 Introduction

Since SARS-CoV-2 infected the human host, its genome has undergone changes leading to the emergence of variants [1]; among the most dangerous, Alpha, Beta, Gamma, and Delta all show changes in the Spike Protein Receptor Binding Domain (RBD). Two trends are prevalent in the evolution of the spike: i) selection towards improved attachment to host cells [2] and higher infectivity; and ii) selection towards evasion of neutralizing antibodies (nAbs) [3], leading to repeat infections [4] and reduced efficacy of vaccinations [5]. The ability to predict the most dangerous spike variants before their natural emergence would have allowed a head-start towards their containment, with substantial societal benefits. The importance of research efforts to anticipate viral evolution has long been established in the scientific community [46] and, in light of recent events, cannot be overstated. Here, among different factors that can drive viral evolution, we will focus on the viral spike’s binding to ACE2 as its natural substrate and to nAbs as one of the host’s countermeasures.

Presently, the main method to reveal potential interactions among spike variants and substrates of clinical relevance is via high-throughput *in vitro* screening of mutants. Two such studies have recently focused on the spike-hACE2 [6] and spike-antibody [7] interactions. These screenings are highly effective at gathering information on a vast chemical space. However, they provide aggregated data, with no direct focus on the mechanisms that make a mutated protein more, or less, dangerous.

To gain mechanistic insight, computational modeling of molecule-molecule interactions can complement wet-bench techniques. Models of inter-molecular interactions have been employed for small molecules (about a hundred atoms) in drug discovery and for protein-protein interactions [8, 9]. Modeling larger molecules remains computationally challenging; nevertheless, several *in silico* approaches, with different sets of assumptions and simplifications, have been successful. Molecular docking uses geometrical constraints to assess how two objects interact with each other [10–12]. Geometry is the main variable in molecular docking, making the method relatively fast and apt at surveying, for instance, small-molecule candidates in drug discovery. Alternatively, force-fields can be employed (FFs), provided an adequate parameterization is found for the system under investigation [18]. FFs employment in biomolecular modeling has over the years consistently brought remarkable results [19, 20]. Hybrid quantum mechanics/molecular mechanics (QM/MM) methods are also common in describing enzyme-substrate systems [13], and can be applied successfully to the characterization of SARS-CoV-2 [15, 16]. QM/MM uses accurate quantum mechanical (QM) simulations for a small fraction of the system, leaving the remaining regions to be modeled with a less computationally demanding MM simulation, driven by FFs. Determining the appropriate QM region for a hybrid model is far from trivial, especially when the sites of interactions on each molecule are unknown [17].

Different computational approaches have focused on the SARS-CoV-2 spike, and the scientific community is actively working on the subject. For instance, coarse-grained modeling was employed to characterize binding to ACE2 and antibodies [40, 41], molecular docking to predict molecular inhibitors [14, 38, 39], molecular dynamics to design peptides against the spike [42], and normal mode analysis to explore conformational states [47]. QM modeling using Density Functional Theory has been employed to analyze functional domains of the Spike protein [44] and to simulate the spike electronic structure to find hydrogen bonds determining its interactions. [43].

A computational method able to perform *ab initio* QM simulations to model interactors across the whole spike structure can complement previous approaches. By focusing on the SARS-CoV-2 spike interaction with select substrates relevant to viral fitness, such as host cell receptor ACE2 and nAbs, we characterize elements of the spike’s evolutionary trajectory and offer predictions on how it will progress. Only recently [21, 34] has the progress in computing capabilities enabled QM simulations of molecular interactions which include tens of thousands of atoms, capturing biological processes involving several hundreds of amino acids. In turn, the SARS-CoV-2 pandemic has recently and rapidly made available a plethora of state-of-the-art biochemical and structural data, facilitating a full QM representation of specific molecular systems. Notably, in the recent years, several full QM calculations of biological macro-molecules of sizes large enough to represent relevant SARS-CoV-2 processes have been published (see e.g. [21] for some examples). Here, we perform *ab initio* QM modeling of one such case: the binding of the SARS-CoV-2 spike RBD with select substrates.

The QM model highlights the chemical hotspots of the viral spike and its interacting protein; we define these hotspots as amino acids with significant energetic contribution to the inter-molecular interactions. The model also reveals the nature of such contributions: chemical/short ranged or electrostatic/long ranged. This mechanistic characterization highlights how mutations in the spike’s primary structure affect binding. Starting from a set of fully atomistic 3D structural models, we employ the BigDFT computer program [22], based on an *ab initio* Density Functional Theory approach, to simulate large molecules with a computational cost manageable on modern supercomputers. Here, BigDFT is applied to the analysis of a set of known cases of viral adaptation, namely: i) the role of the E484K mutation in the evasion of nAbs C121 and C144 [5]; ii) the role of the spike residue 484 in SARS-CoV-2’s affinity for the human ACE2 receptor (hACE2); iii) the spike interaction with the bat host *Rhinolophus macrotis* ACE2 (macACE2); and iv) the Delta variant spike interaction with hACE2. Finally, we use the model to predict the effect of a hypothetical mutation of the SARS-CoV-2 Delta variant spike RBD in the binding to human cells.

## 2 Methods

### Computational approach

Our study is performed via a full Quantum Mechanical (QM) model, as implemented in the BigDFT computer program suite [31]. The approach employs the formalism of Daubechies wavelets to express the electronic structure of the assemblies in the framework of the Kohn-Sham (KS) formalism of Density Functional Theory (DFT) [22]. The electronic structure is expressed by both the density matrix and the Hamiltonian operator in an underlying basis set of support functions – a set of localized functions adapted to the chemical environment of the system. Such functions are expressed in Daubechies wavelets, typically using one to four support functions per atom as the basis set. The electronic density matrices, as well as the Hamiltonian expressed in the BigDFT basis set, are analyzed to provide quantum observables of the systems. The code provides efficient and accurate QM results for full systems of large sizes, delivering excellent performance on massively parallel supercomputers. In the present study, we employ the PBE approximation corrected by dispersion D3 correction terms [36]. Each of the calculations presented here requires about 2 h of wall-time on 32 compute nodes of the IRENE-Rome supercomputer, at the TGCC Supercomputing center in Saclay (Paris, France). A similar approach has been previously used, in conjunction with the other atomistic techniques described in the introduction, to investigate the interaction patterns of the SARS-CoV-2 main protease with natural peptidic substrates and to design peptide inhibitors tested *in vitro* [25].

### Procedure

Starting from a representative 3D model of the molecules as our input, we calculate the system’s electronic structure, from which we extract various quantities. In particular, we draw a contact map to identify relevant chemical interactions between the spike RBD and the various interactors considered in this study. The strength of the inter-residue interaction is quantified by the Fragment Bond Order (FBO) [24], calculated using the electronic structure of the system in proximity of a given residue. Such an approach has been previously described in detail [22] and is summarized in Table 1. We use the FBO to identify the interface residues, defined as the amino acids of the counter-ligand that have a non-negligible value, above a set threshold of the FBO, with the ligand. In contrast to a simple geometrical indicator like the RBD-ligand distance, the FBO provides a metric that enables a non-empirical identification of steric hot-spot interactions. We here identify as chemical hot-spot interface residues the amino acids which exhibit a FBO value with the ligand larger than 7 · 10^−3^. Such threshold value has been chosen from a comparison between the hydrogen bonding interaction network of the SARS-CoV-2 main protease with its natural peptidic substrates, derived from traditional FF analysis, and the equivalent FBO network [21].

**Table 1.**
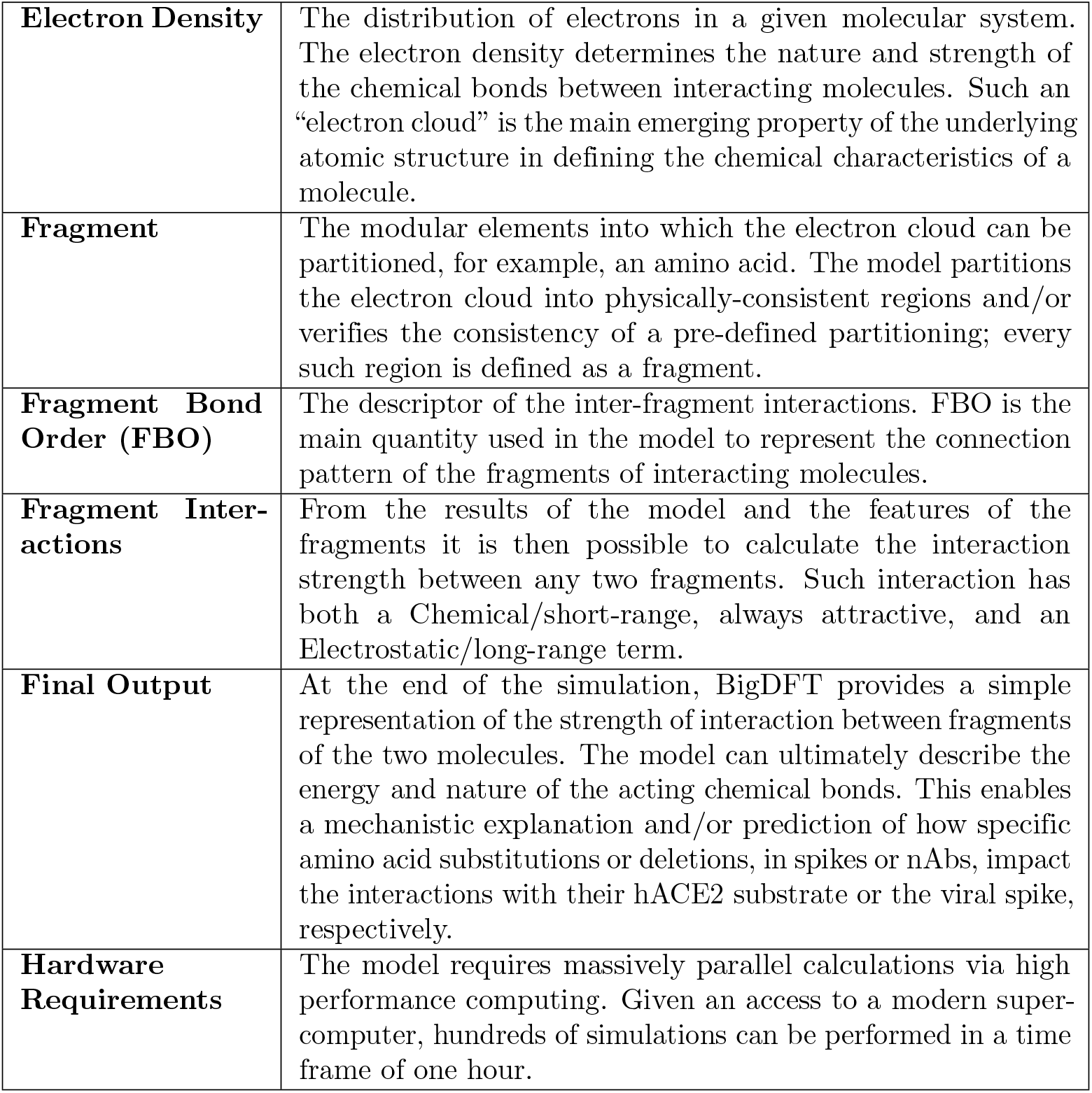
Prospectus of the main concepts and quantities constituting the model. All the elements here discussed are general and therefore applicable, without previous parameterization, to any given set of atoms for which atomistic structural representations are available.

|  |  |
| --- | --- |
| **Electron Density** | The distribution of electrons in a given molecular system. The electron density determines the nature and strength of the chemical bonds between interacting molecules. Such an “electron cloud” is the main emerging property of the underlying atomic structure in defining the chemical characteristics of a molecule. |
| **Fragment** | The modular elements into which the electron cloud can be partitioned, for example, an amino acid. The model partitions the electron cloud into physically-consistent regions and/or verifies the consistency of a pre-defined partitioning; every such region is defined as a fragment. |
| **Fragment Bond Order (FBO)** | The descriptor of the inter-fragment interactions. FBO is the main quantity used in the model to represent the connection pattern of the fragments of interacting molecules. |
| **Fragment Interactions** | From the results of the model and the features of the fragments it is then possible to calculate the interaction strength between any two fragments. Such interaction has both a Chemical/short-range, always attractive, and an Electrostatic/long-range term. |
| **Final Output** | At the end of the simulation, BigDFT provides a simple representation of the strength of interaction between fragments of the two molecules. The model can ultimately describe the energy and nature of the acting chemical bonds. This enables a mechanistic explanation and/or prediction of how specific amino acid substitutions or deletions, in spikes or nAbs, impact the interactions with their hACE2 substrate or the viral spike, respectively. |
| **Hardware Requirements** | The model requires massively parallel calculations via high performance computing. Given an access to a modern super-computer, hundreds of simulations can be performed in a time frame of one hour. |

Once the chemical connection between amino acids is identified, we assign to each of the residues its contribution to the binding interaction between the two subsystems. Such interaction terms can be calculated from the output of the DFT code, and can be split in two parts. The first is a electrostatic attraction/repulsion term, defined from the electron distributions of each of the fragments, which has a long-range character (even when they are far apart two fragments may still interact). The remaining term, which can only be attractive, is provided by the chemical binding between two fragments, and is non-zero only if the electronic clouds of the fragments superimpose (short-range). This term is correlated to the FBO strength, and we identify it as the chemical interaction. By including long-range electrostatic terms, the decomposition enables us to single out relevant residues not necessarily residing at the interface. In this way, the model provides an *ab initio* representation of the RBD-ligand interactions as the final output.

### Crystal structures and generation of virtual structures for mutants

Crystallographic structures are obtained from the RCSB database [26](PDB entries 6M0J (hACE2), 7K8X (nAb C121), 7K90 (nAb C144), and 7C8J (macACE2)). Protonation of histidines and other titratable residues is assigned a pH of 7, based on the PDBFixer tool in OpenMM [27]. Virtual structures are generated by the imposition of point mutations of the original structure via the same tools. Structure relaxations are performed by optimizing the crystal geometry with the OpenMM package using the AMBER FF14SB force field [28]. While such optimized structures do not represent the full panorama of conformations that might exist at finite temperature, the resulting structures are interpreted as one plausible representative among the possible conformations of the system.

## 3 Results

We focus our analysis on the impact of the E484K mutation on antibody evasion and receptor binding. Prior experimental data have shown that antibodies C144 and C121 are evaded completely by spike variants presenting the E484K mutation in the RBD [5]. E484K is a typical signature mutation of the RBD of the Gamma and Beta strands. We test our QM model as an agnostic predictor to explain existing biological data, and to characterize the underlying chemical interaction of the nAbs with both the original Wuhan spike and the E484K-mutated one.

In Fig. 1, for each amino acid of the primary structure, we represent its contribution to the binding energy, which can be attractive/stabilizing or repulsive/de-stabilizing. We also highlight FBO-interface residues (Fig. 1 yellow bars) as well as those close to the geometric interface. In the forthcoming section we will use FBO to draw an interaction network of the interface to detail the chemical interactions among residues and their role as stabilizing or destabilizing agents. Details of the procedure are provided in the supplementary A.4.

**Fig 1.**
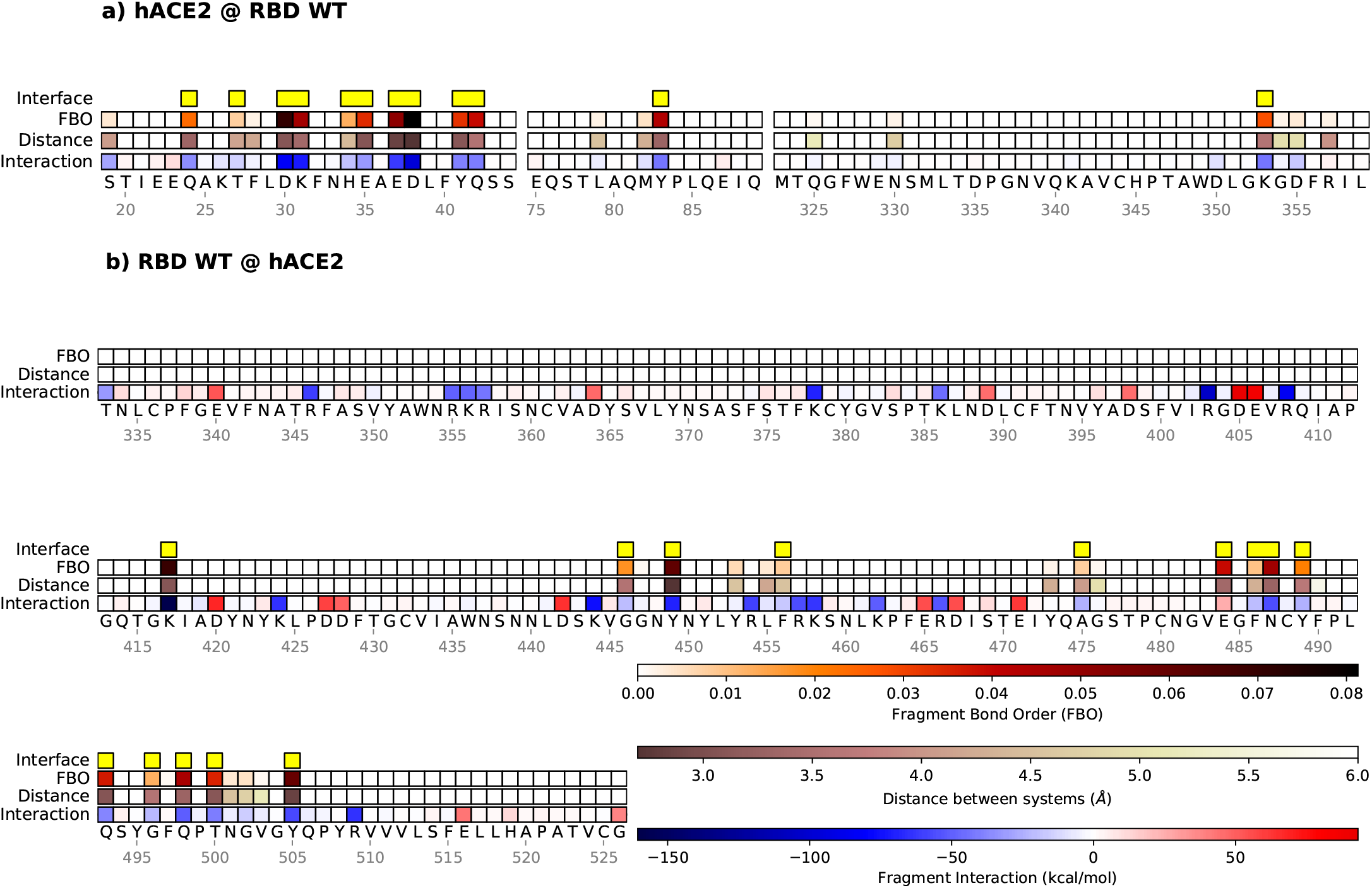
Mechanistic characterization of the binding between Wuhan strain’s spike and hACE2. Data are plotted on the sequence of hACE2 (panel a) and the spike RBD (panel b). Letters represent single amino acid residues; yellow bars indicate interface residues, identified with the FBO threshold. The “FBO” row represents Fragment Bond Order values, the “Distance” row represents the distance of a residue to the nearest atom of its ligand, and the “Interaction” row shows the chemical/electrostatic force as attractive (blue) or repulsive (red), with darker colors indicating stronger effects.

### 3.1 *Ab initio* simulation shows how nAb C121 loses binding to the E484K mutated spike

The first step in the analysis identifies the hotspots between the RBD and the nAbs of the Wuhan spike interaction (Fig. 2). Residue E484 emerges as the main spike interactor with the nAb C121. Other relevant sites of interaction are residues K444, Y449, F486, Y489, and Q493. On the C121 side, residues Y33, S55, and S75 are identified as pivotal for the Wuhan spike binding. The model estimates that among all the residues contributing to the interaction, the individual contribution of E484 amounts to roughly 50% of the total. The interaction network of the assembly (see the second part of Fig. 2) completes the characterization of E484 by showing its coordinated binding to elements on the C121 structure; namely, residues Y33 and S55. By imposing the E484K mutation, we observe a rearrangement of the interaction network and a substantially lower binding energy between the spike and the antibody. Specifically, E484K reduces the connectivity at the 484 residue in the interaction network, and modifies the interactions on the C121 side towards decreased stability. Only the S52 residue takes advantage of the mutation, but its stabilizing contribution is insufficient to counterbalance the loss of attraction at other residues. Overall, once the mutation is applied, we observe a substantial decrease of about 25% of total binding energy, largely attributable to loss of chemical interaction. The model concludes, with no *a priori* information, that the E484 residue is the essential actor in the binding by nAb C121, and that a targeted point mutation will substantially affect said binding. The analysis of C144 nAb shows similar results. Moreover, C144 undergoes a substantial rearrangement of its interaction network in response to the mutation, arguably a consequence of the original higher connectivity of the residue E484 in the binding, compared to the C121 case: five coordinated residues (Y51, S52, G53, G54 and S55) instead of two (Y33 and S55) (Supplementary Fig. A.1). Interestingly, the importance of E484 also appeared in previous results by Andreano *et al.* in which E484 mutants arise under the selective pressure of nAbs [33].

**Fig 2.**
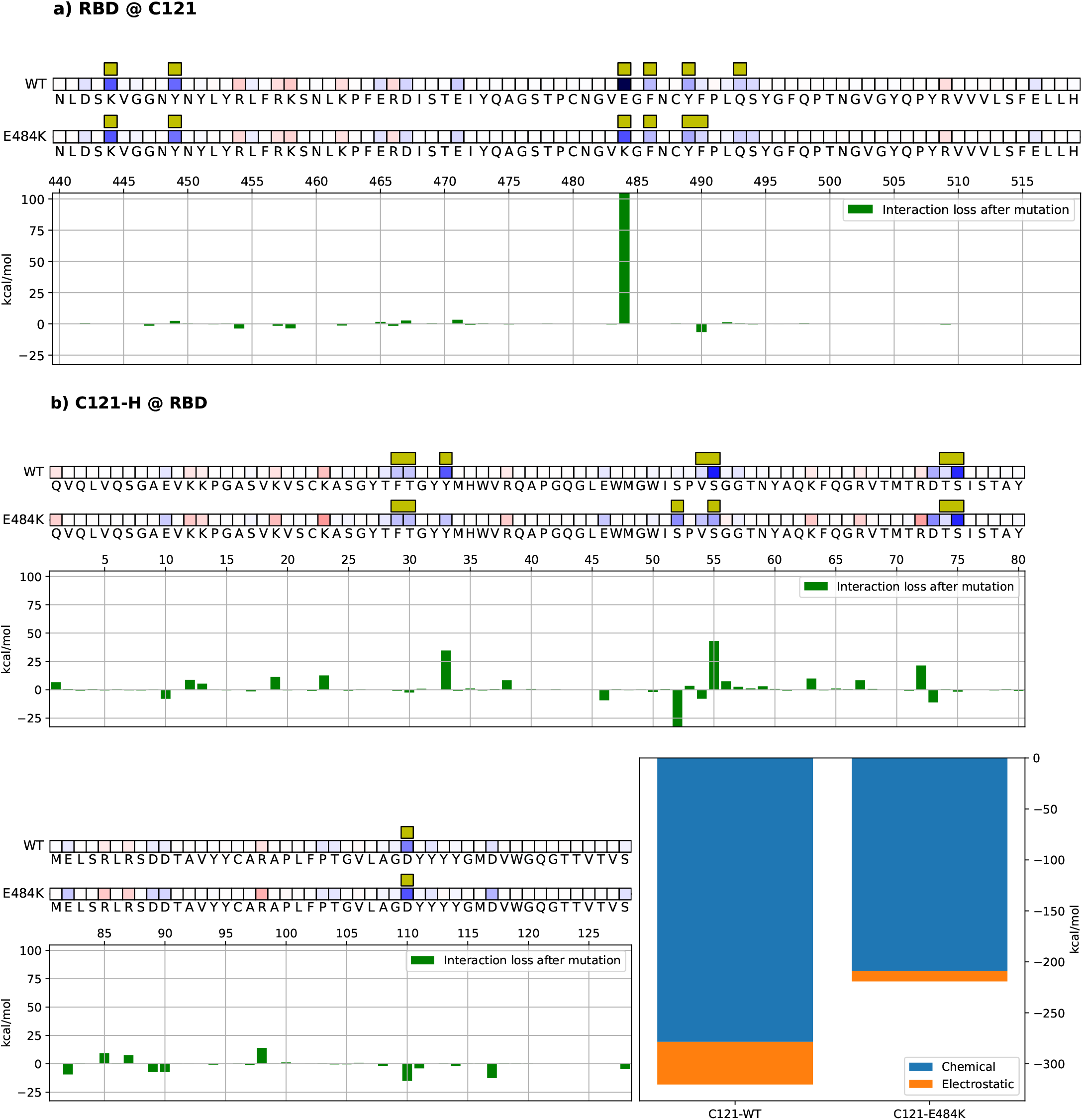
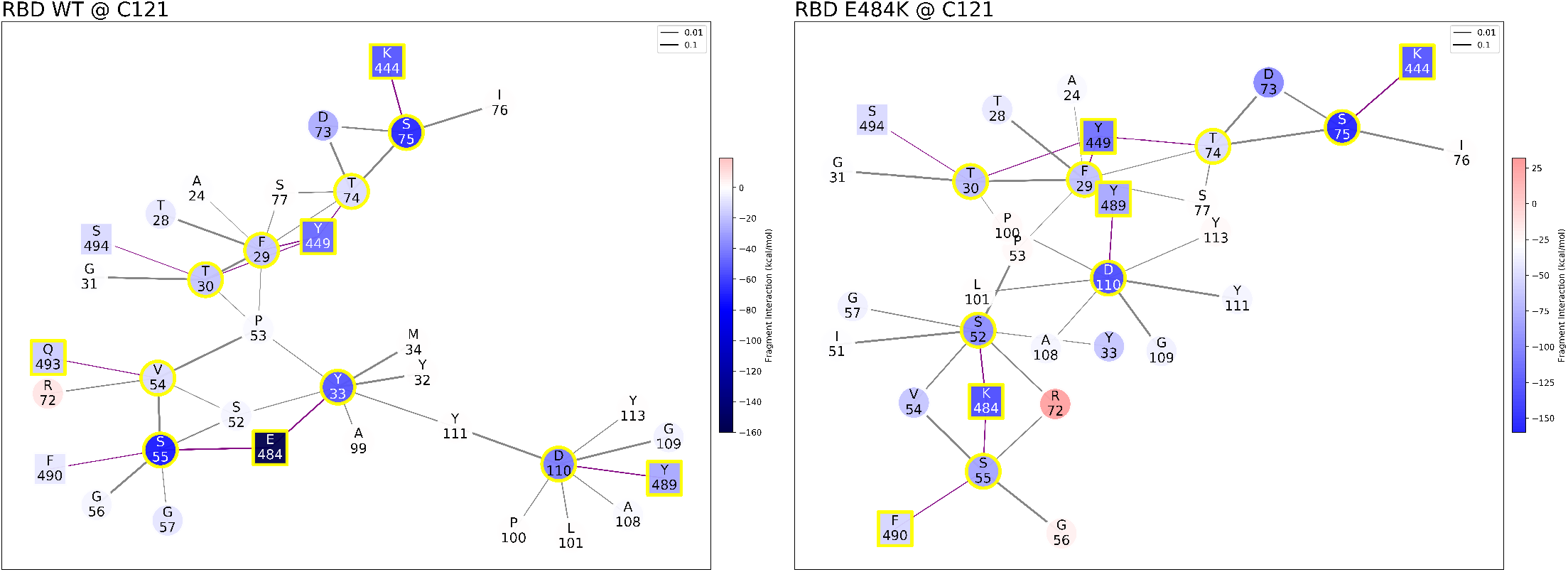
Mechanistic characterization of C121 binding to the Wuhan strain spike protein, and energetic changes as a result of the E484K spike mutation. Data are plotted on the spike primary structure (panel a) and on C121 Heavy-Chain (panel b) considering the different bindings via the Wuhan spike (WT) and the mutated one (E484K). Amino acids are represented by the letters, and numbered on the histogram’s horizontal axis. Histograms underneath the sequences represent the relative change in binding energy of the second row relative to the first one (Wuhan type strand). The bottom right histograms represent the overall binding energy of C121 with the Wuhan spike (left) and the mutated one (right) and its characterization as chemical or electrostatic. The row above each sequence shows the chemical/electrostatic force as attractive (blue) or repulsive (red), with darker colors indicating stronger effects. Interaction networks with C121 nAbs. Bonds are purple when inter-molecular or black when intra-molecular, and their thickness is related to the strength of the FBO between residues. Graph nodes are represented in red (repulsive) and blue (attractive) based on their effect on their counterpart. Residues at the binding interface are highlighted by a yellow outline.

### 3.2 *Ab initio* simulation identifies the spike E484 residue as the weak link in the binding to the host receptor hACE2

The FBO identifies the chemical/short ranged interactions, bringing out the hotspots of the RBD-hACE2 system (Fig. 3). On the hACE2 side (Fig. 3, panel a), Q24, T27, D30, K31, H34, E35, E37, D38, Y41, Q42, Y83, and K353 stand out, in agreement with known data [30]. On the spike side (Fig. 3, panel b), a more diverse layout emerges, on and off the interface, with several residues displaying repulsive interaction. However, residue E484 shows the unique trait of being contextually repulsive and at the interface with hACE2, via a short range interaction with the K31 residue. This implies, as the chemical interaction is intrinsically stabilizing, that another residue in the vicinity hinders its stabilization with an electrostatic repulsion. Therefore, in the Wuhan type structure, E484 is actually destabilizing the binding to hACE2. From this analysis, we conclude that the Wuhan spike RBD harbors a sub-optimal residue for hACE2 binding.

**Fig 3.**
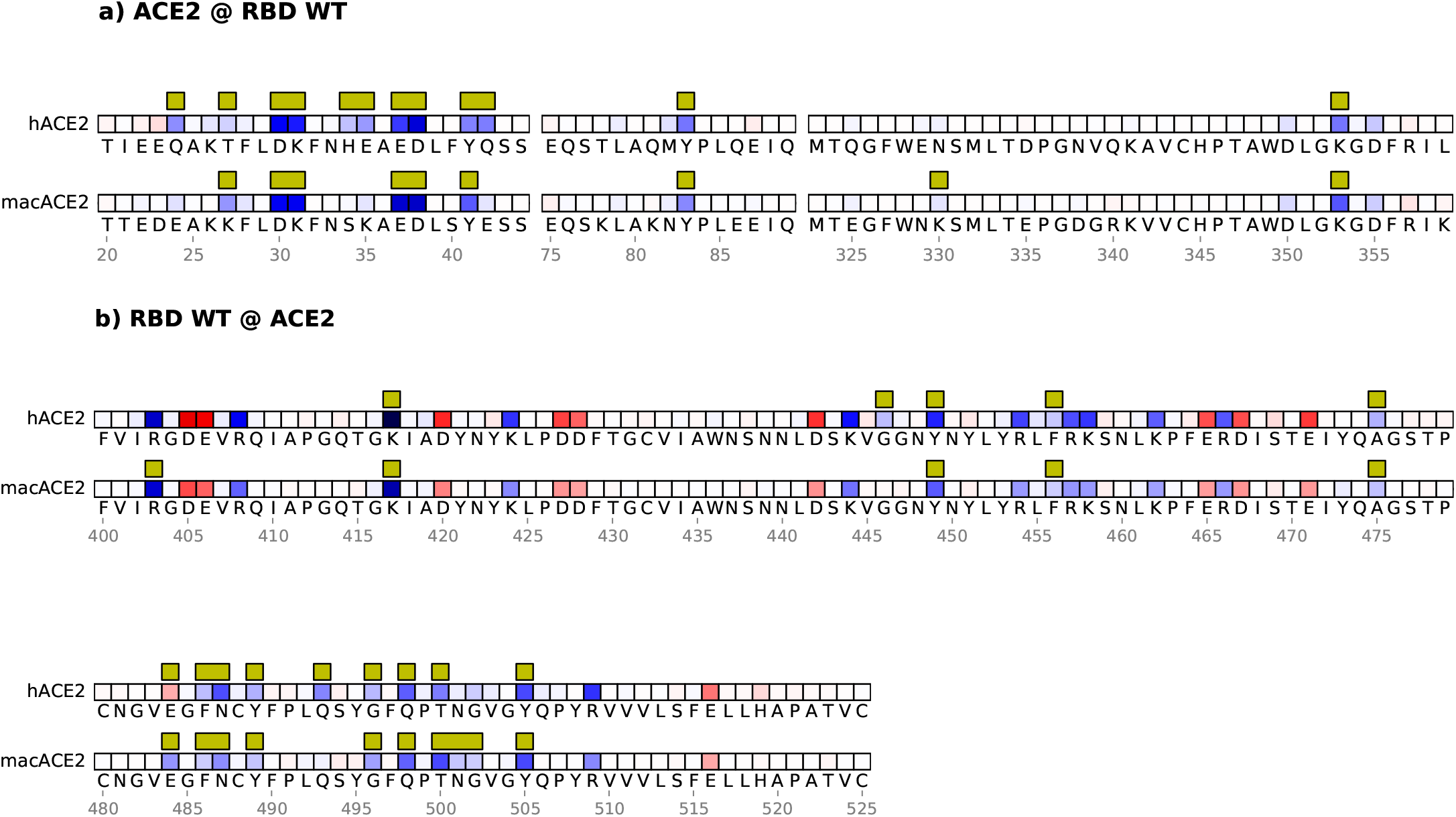
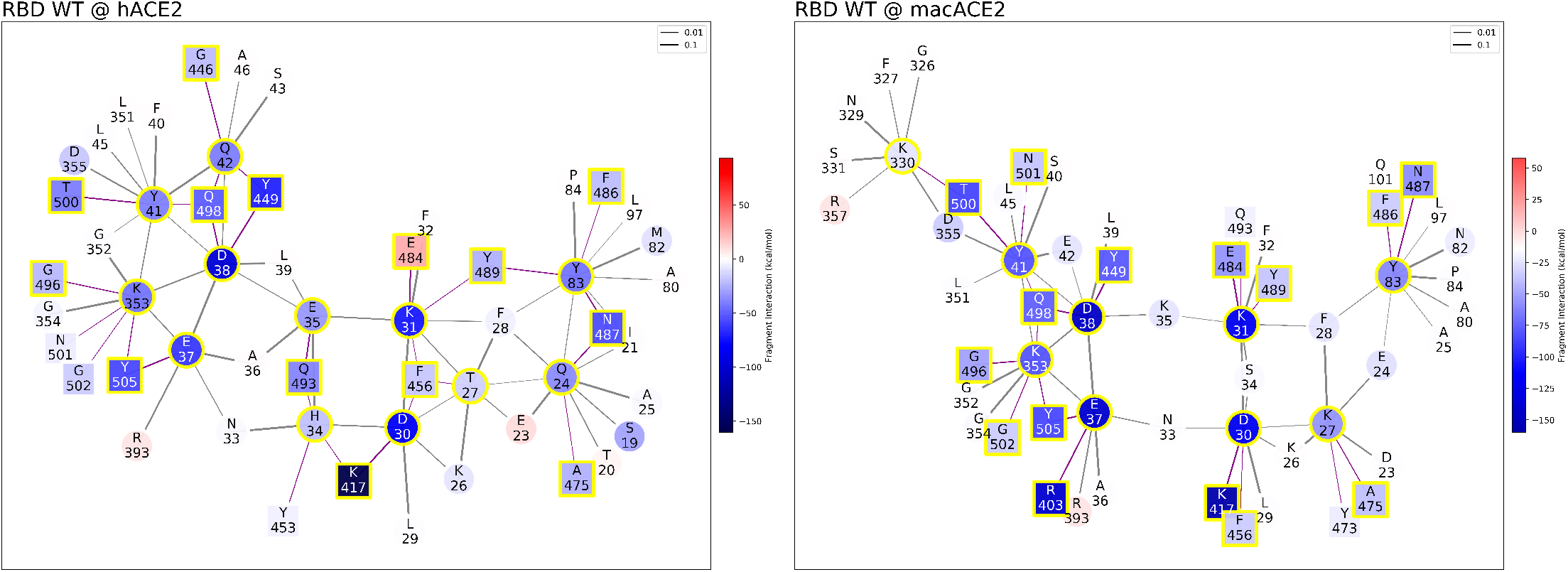
Mechanistic characterization of the Wuhan spike binding to the human ACE2 (hACE2) and *R. macrotis* ACE2 (macACE2). Data are plotted on the ACE2 primary structure (a), and on the Wuhan spike RBD (b), when binding to the human (hACE2) and the bat (macACE2) receptor. Amino acid residues are labeled with letters and numbered. Interface residues are highlighted with a yellow bar, red tiles are repulsive residues, and blue tiles are attractive residues; see the rest of the figure for energy scales. The interaction networks represent the hACE2-spike system on the left, and macACE2-spike on the right; circles are ACE2 residues, squares are spike residues. Interface residues are highlighted with a yellow bar, red tiles are repulsive residues, and blue tiles are attractive residues. Bonds are purple when inter-molecular or black when intra-molecular.

To better investigate this point, we test the model on the available 3D crystal structure of the human homologous ACE2 receptor in *Rhinolophus macrotis*, a host species with arguably a more adapted SARS-CoV-2 interaction [30]. In this simulation (Fig. 3 panels a and b, second rows), we observe the E484 residue is instrumental to the binding by being strongly attractive to the *R. macrotis* ACE2 (macACE2); notably, in both hACE2 and macACE2, the interactor with E484 is the ACE2 residue K31. This means that the macACE2 sequence has residues, proximal to the K31 hotspot, that exert an attractive electrostatic force on E484. A closer inspection of the two sequences reveals that this attractive force comes from the K35 residue, which in hACE2 is replaced by Glutamic Acid. The model therefore highlights a strong contrast between human and bat receptors.

The role of E484 is further confirmed by the analysis of the spike virtual structure with the E484K mutation imposed interacting with hACE2 (Fig. 4): hACE2 binding improves by about 32% ΔE in presence of the mutation (Fig. 4, bottom right histograms), switching its original interacting residue from K31 to E35. Such an interaction, driven by electrostatics, represents a net improvement of the network. Conversely, the same mutation does not affect the spike binding energy to macACE2 in the same position, where the bat receptor hosts a Lysine. In other terms, for macACE2, K484 clearly does not engage K35, and actually disappears from the interface (Supplementary Fig. 4); the resulting interaction network is largely rearranged, and the interface binding energy is not improved by the mutation. Therefore, the model shows an arguably more optimal interaction between macACE2 and WT RBD, possibly the result of a long adaptation by SARS-CoV-2 to *R. macrotis*, whereas in the hACE2 receptor, the E484 residue belongs to a sub-optimal sector of the chemical interface, suggesting that other RBD adaptations in this sector are likely to improve the binding.

**Fig 4.**
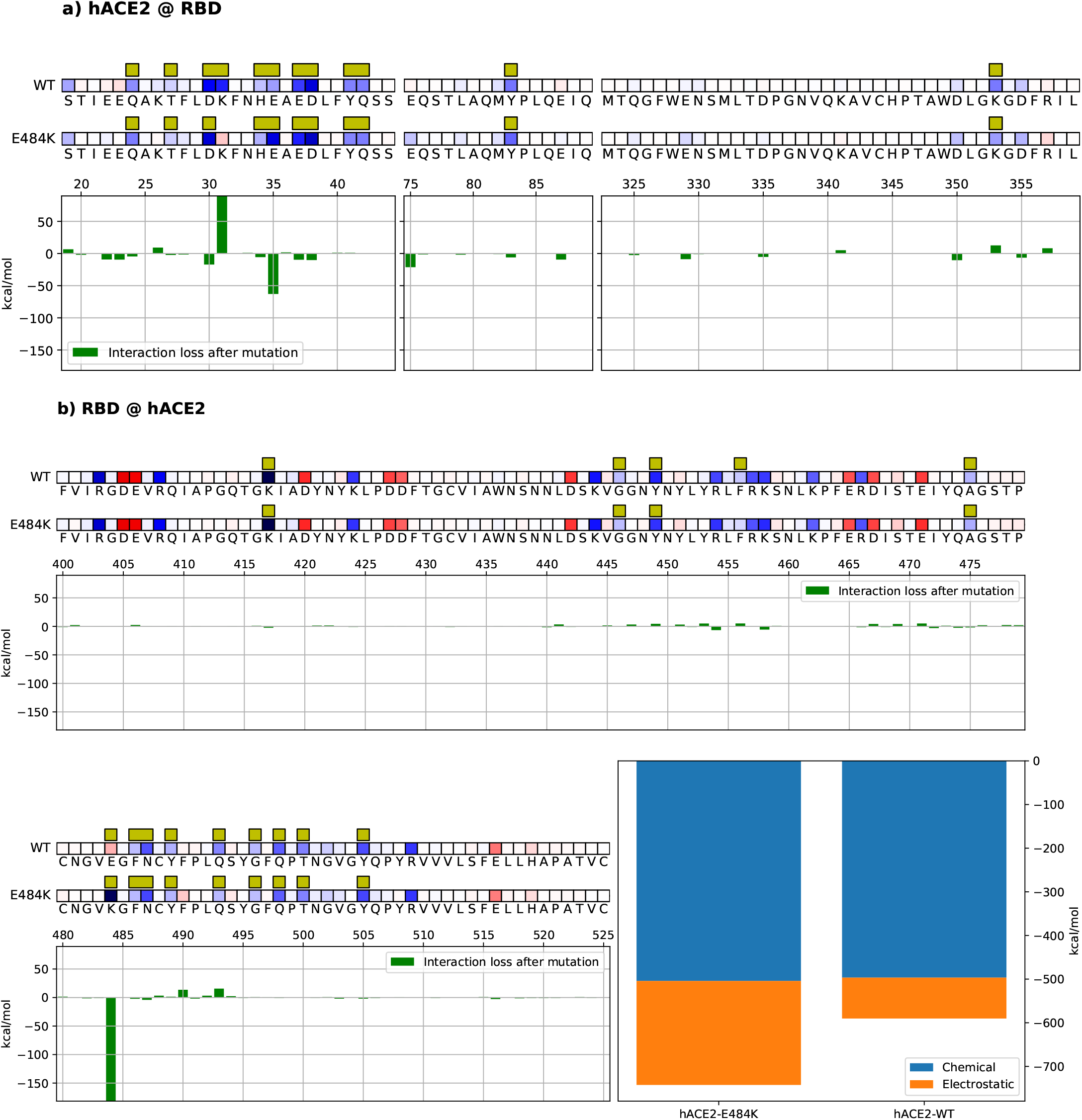
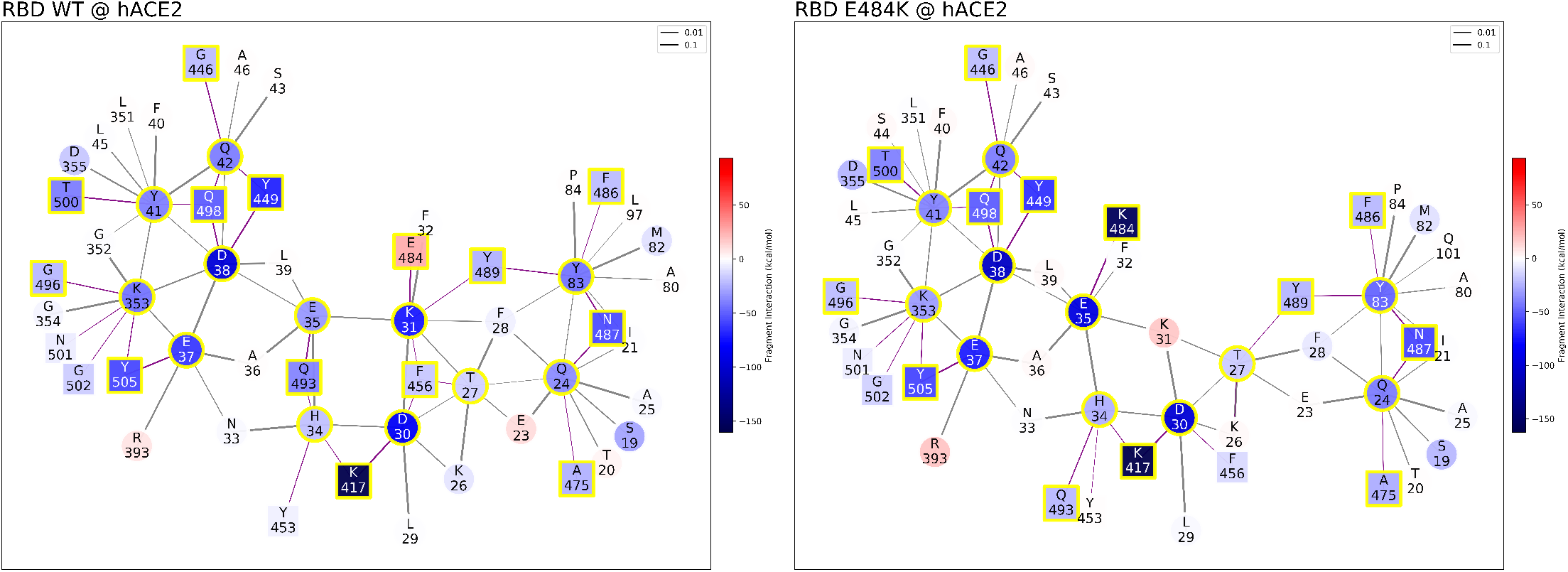
Mechanistic characterization of Wuhan and mutated (E484K) spike binding to hACE2. Data are plotted on hACE2 (panel a) and on the Wuhan spike (panel b) primary structure bound to the Wuhan spike (WT) and the mutated one (E484K). Amino acids are represented by the corresponding letters, and numbered on the histogram’s horizontal axis. Interface residues are highlighted by yellow bars and their overall effect on the other molecule is indicated by red (repulsive) or blue (attractive) tiles. Histograms underneath the sequences show the relative change in binding energy of the second row (E484K mutation) relative to the first one (WT strand), with positive and negative values indicating weaker and stronger binding, respectively. Bottom right histograms represent the overall binding energy of hACE2 with the Wuhan spike versus the mutated one, partitioned into chemical and electrostatic contributions. Interaction networks (Wuhan spike-hACE2 to the right, and mutated spike-hACE2 to the left), including FBO-interface residues and their coordinated interactors. Squares depict spike residues and circles depict hACE2 residues, with red color for repulsive and blue color for attractive energy. Yellow outlines highlight interface residues. Bonds are purple when inter-molecular or black when intra-molecular.

### 3.3 *Ab initio* simulation predicts that the E484K mutation increases binding of Delta spike to hACE2

Presently, the Delta variant’s high infectivity is a major concern in the COVID-19 pandemic. At the time of writing this contribution, an experimentally validated Delta spike-ACE2 3D crystal structure is not publicly available. We thus generate a virtual crystal structure to represent Delta (B.1.617.2) in conjunction with hACE2 by substituting its characterizing RBD mutations (L452R and T478K) into the Wuhan spike crystal structure. Such residue mutations belong to a far-from-interface sector of the RBD (see Fig. 1). Our simulations identify the same FBO interface residues found for the Wuhan strain. However, differently from the other tested interaction networks, a substantial contribution to the overall binding energy of Delta to hACE2 comes from off-interface residues via their long range electrostatic effect on their counterparts, highlighting the relevance of including residues beyond the interface region in the analysis of binding.

Furthermore, when testing the binding of the Delta-hACE2 system after introducing the E484K mutation, the simulation shows that E484K is compatible with the present Delta variant and further strengthens the overall binding to hACE2. Such an *in-silico* variant, at present solely based on theoretical grounds, displays a stronger binding to hACE2 than either E484K and Delta variants individually (Fig. 5).

**Fig 5.**
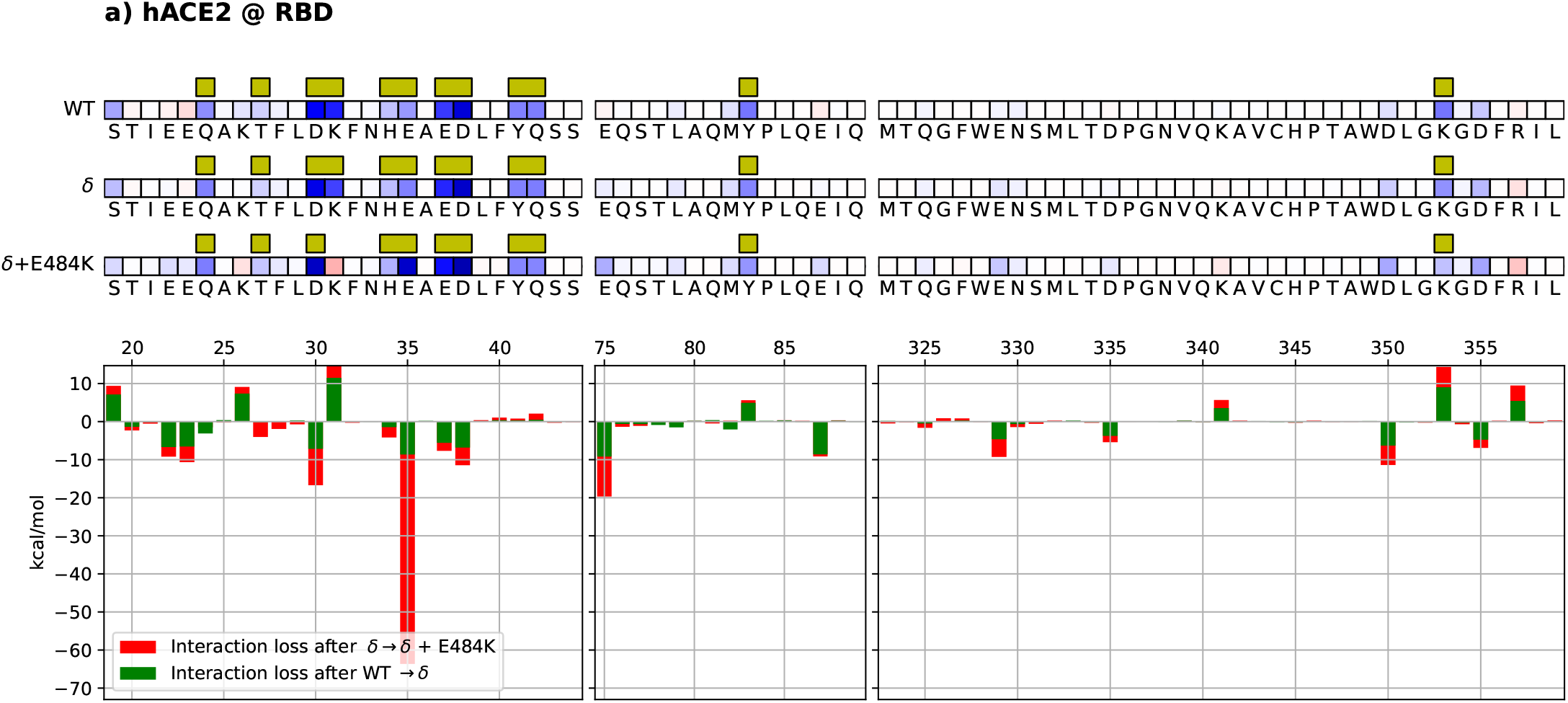
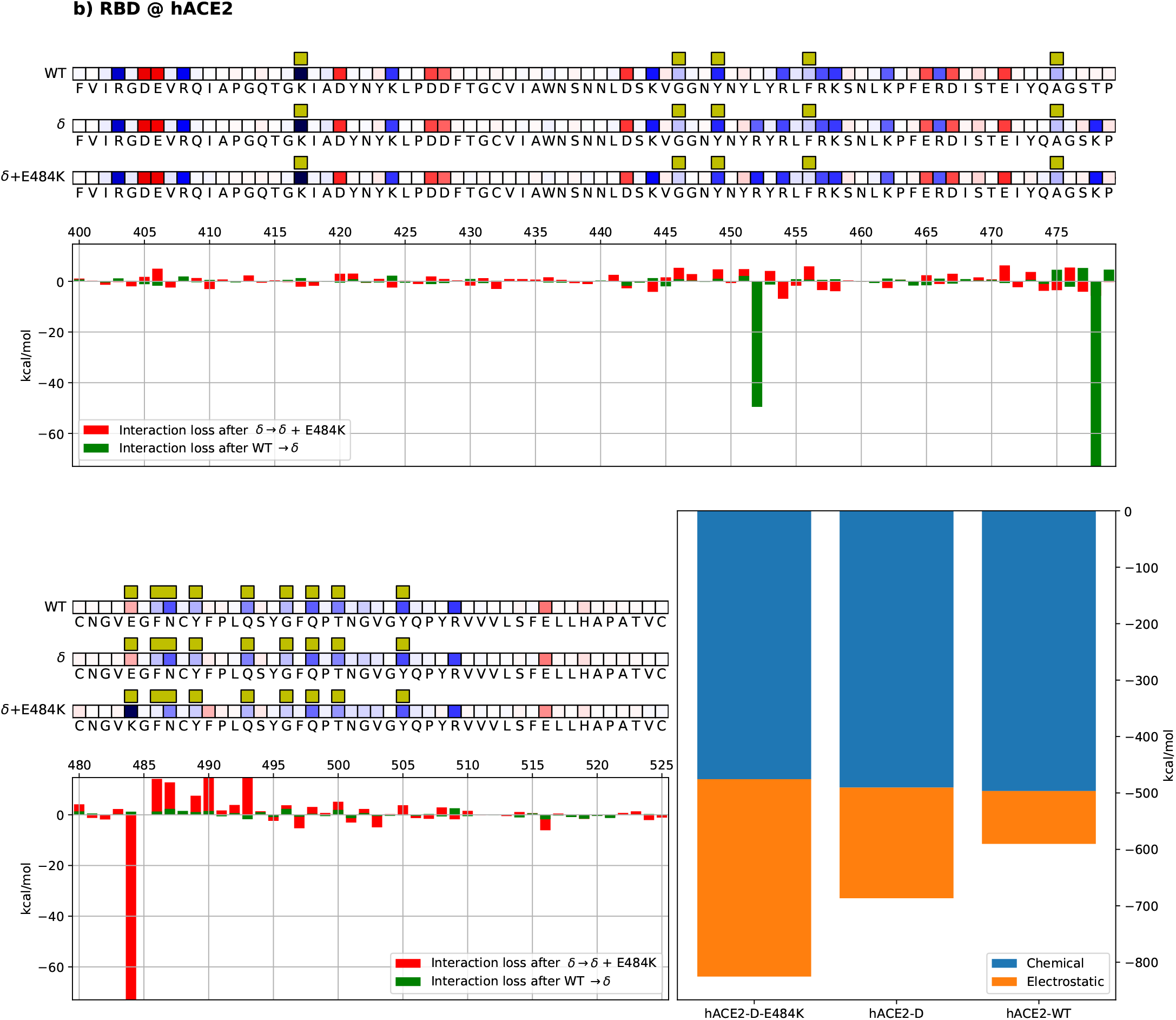
Mechanistic characterization of spike-hACE2 binding suggests that Delta+E484K spike has stronger hACE2 binding that the Delta variant. Data are plotted on hACE2 (panel a) and viral spike (panel b) primary structure bound to the Wuhan spike (WT), Delta spike (*δ*), and Delta + 484K spike (*δ* + 484K). Amino acids are represented by the corresponding letters, and numbered on the histogram’s horizontal axis. Interface residues are highlighted by yellow bars and their overall effect on the other molecule is indicated by red (repulsive) and blue (attractive) squares (energy scale is identical to the one employed in the other figures). Histograms underneath the sequences show the relative change in binding energy (green: Delta compared to Wuhan; red: Delta+E484K compared to Delta). Bottom right histograms represent the overall binding energy of hACE2 with the Wuhan, Delta, and Delta+E484K strains, partitioned into chemical or electrostatic contributions.

## 4 Discussion

The characterization of inter-protein interactions is central to the mechanistic understanding of biological phenomena. The properties of a protein are only partially defined by the residues that constitute its active site. A mechanistic description of protein-protein interactions can benefit from a full-sequence QM characterization. We use the BigDFT code [22, 31] to implement a *ab initio* QM simulation of the electronic properties of a given set of atoms as large as a full protein-protein system. Through this model, we decompose the interaction between two biological macro-molecules, spike and receptor/antibody, into the individual energetic contributions of each of the amino acid residues involved. Additionally, the model characterizes the nature of these contributions into two main categories: (1) short-range/chemical and (2) long-range/electrostatic. Ultimately, we infer a network of interactions in which every node is a single amino acid belonging to one of the two molecules; the network is based on the electron cloud surrounding the protein-protein system, itself an emergent property of the structural arrangement of the system’s atoms.

In this work, we examine how the viral spike interacts with ACE2 as its natural receptor, and with nAbs C121 and C144. We demonstrate that a QM model, assessing the interactions among the residues of an inter-molecular biological system, enables mechanistic insight into how SARS-CoV-2 interacts with its host.

The model identifies the E484 residue as the only interface element hindering the binding between the Wuhan strain and hACE2, *de facto* making it the most evident weak link of the Wuhan spike binding to the human host. The E484K mutation is shown by the model as a direct solution to this hindrance by improving binding to hACE2, and presumably constituting an evolutionary advantage which led to its emergence among several successful variants. Interestingly, the *ab initio* model also shows that the E484 residue does not destabilize the interaction between the Wuhan viral spike and the bat receptor macACE2 from *Rhinolophus macrotis*. We interpret this result as an indication that the Wuhan strain is adapted to a bat-like ACE2, and the rise of E484K variants is indeed part of the viral adaptation specific to the human host.

We find that the *ab initio* model correctly predicts the loss of interaction between the SARS-CoV-2 spike and nAbs C121 and C144, once the E484K mutation is imposed on the spike of the Wuhan strain. The RBD residue E484 emerges as the main and fundamental spike fragment enabling the binding event, and therefore neutralization. These data suggest that nAbs challenging the spike at E484, the very residue that most hinders hACE2 interaction, have provided selective pressure for the virus to find alternatives to the original phenotype.

The binding energies provided by our model can be compared against quantities obtained from experimental databases of the libraries of the interaction between spike RBD mutants and hACE2 mutants available in the literature [6, 7]. While the two quantities are not directly comparable, as computational studies of protein-protein affinity requires in-depth analysis of structural and thermodynamic contributions [48], results from both datasets are in general accordance (Supplementary Fig. A.3). We plan a detailed study of the comparison between such datasets particularly focused on outliers. We argue that *ab initio* simulations have achieved the maturity to inform the exploration of the available chemical space via single-point mutations.

By analyzing the competition between short- and long- range interaction contributions, we have shown that the charge-shift E484K mutation conferred a substantial binding energy increase (about 30%) to the interaction with hACE2, compared to the Wuhan strain. On the RBD side, the model also highlights how the effect of E484K is focused on the 484 position, with limited off-target repercussions for the spike’s binding (Supplementary Fig. 4). We argue that this trait qualifies the E484K mutation as highly “RBD-modular”, therefore easy to impose on an already functional spike structure, its contribution to the binding being largely long-range/electrostatic, therefore less dependent of a specific steric conformation.

Our *ab initio* simulation is motivated by the available empirical data in identifying the E484K variant as a particularly dangerous evolutionary outcome, based on increased SARS-CoV-2 infectivity and antibody evasion. From this last standpoint, we examined the potential impact of the E484K mutation improving spike-hACE2 binding in the background of the recently spread Delta variant. Our model suggests that E484K affects spike-hACE2 and spike-nAb binding in a modular fashion. Thus, if acquired by the Delta strain, E484K constitutes a likely future threat, to the extent that receptor binding can contribute to an increase in infectivity. We acknowledge that infectivity is a multi-factor process, and receptor binding is only one of the factors involved.

We view our investigation in two lights: on one side, the disruptive nature of the COVID-19 pandemic and its extension because of the emerging variants of SARS-CoV-2. This is a primary motivation to increase our readiness to handle currently emerging variants and anticipate future ones. On the other side, the abundance of data for SARS-CoV-2 has provided an opportunity to test and validate the potentials and limitations of *ab initio* modeling. This, in conjunction with the maturity of large-scale quantum mechanical calculations, represents a unique opportunity to employ full QM calculations to uncover the interaction mechanisms which would be difficult or impossible to investigate otherwise. We show that *ab initio* modeling provides insights useful for comparison with experimental data, supporting its capability to offer predictive power for inter-molecular interactions of biological relevance. Research paradigms on large biomolecules investigation, in various domains, can now effectively include QM data.

### Model limitations

The model, despite its first-principles origin, is based on assumptions. First, the quantities we are investigating, while representing the binding energy, do not directly manifest experimental results. A closely-related quantity may be the off-rate dissociation constant, that can be inversely proportional to the structures’ stability. However, to correctly evaluate such terms, the systems’ free energies must be considered, which would require dynamic structural investigations. Additionally, the mutated structures are based on reference structures whose 3D conformation may in principle be altered by a given mutation. Therefore, in the absence of confirmed crystal structures, these structures can only be interpreted as virtual best guesses. The evaluation of mutations at the interface, particularly mutations which are associated to chemical interactions, should be treated with care, considering the interplay between the binding and the actual position of the residue at the interface. In addition, for electrostatic interactions, solvation effects may affect the relevance of a charged residue; also Glycation has been shown to be relevant in the binding of the Spike protein [37][WD: Comment:reference formatting is off]. However, these mechanisms are independent of the QM treatment, and the investigation of such contributions can be effectively performed by coupling our model to other mechanistic treatments, such as polarizable force fields or advanced docking techniques. In addition, we here employ a common, well-established DFT approximation (PBE+D3), which already provides useful information for coarse-grained quantities and trends [21, 35], and aptly fits the aim of intensive, high-throughput simulations of several structures, especially in their relaxed positions [45]. Yet, we leave out arguments about the choice of other *ab initio* levels of theory, which may shed light on more quantitative aspects related to processes beyond the ground-state: reaction coordinates, activation barriers, etc. For the study presented here, considering that experimental quantities are indirectly related to those data, we believe that the method employed represents an ideal compromise between accuracy of the results and affordable modelling.

We argue that our approach, albeit simplified by the assumptions and limitations described above, can predict and characterize potential antibody escape routes of SARS-CoV-2 and, being unbiased and agnostic, is ready to be applied to other biological systems.

## A Supporting information

### A.1 *Ab initio* simulation shows how nAb C144 loses binding to the E484K mutated spike

**Fig A.1.**
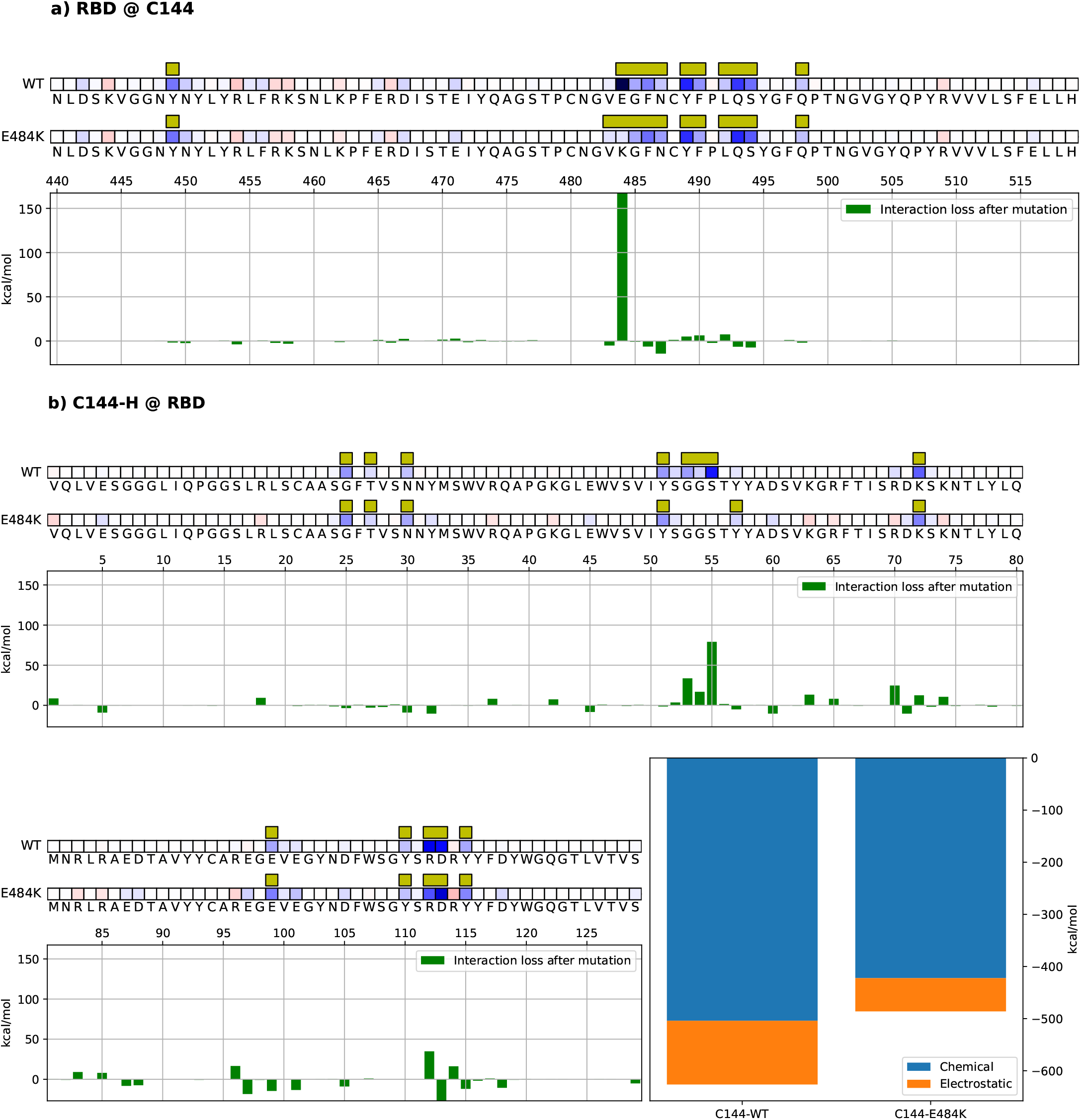
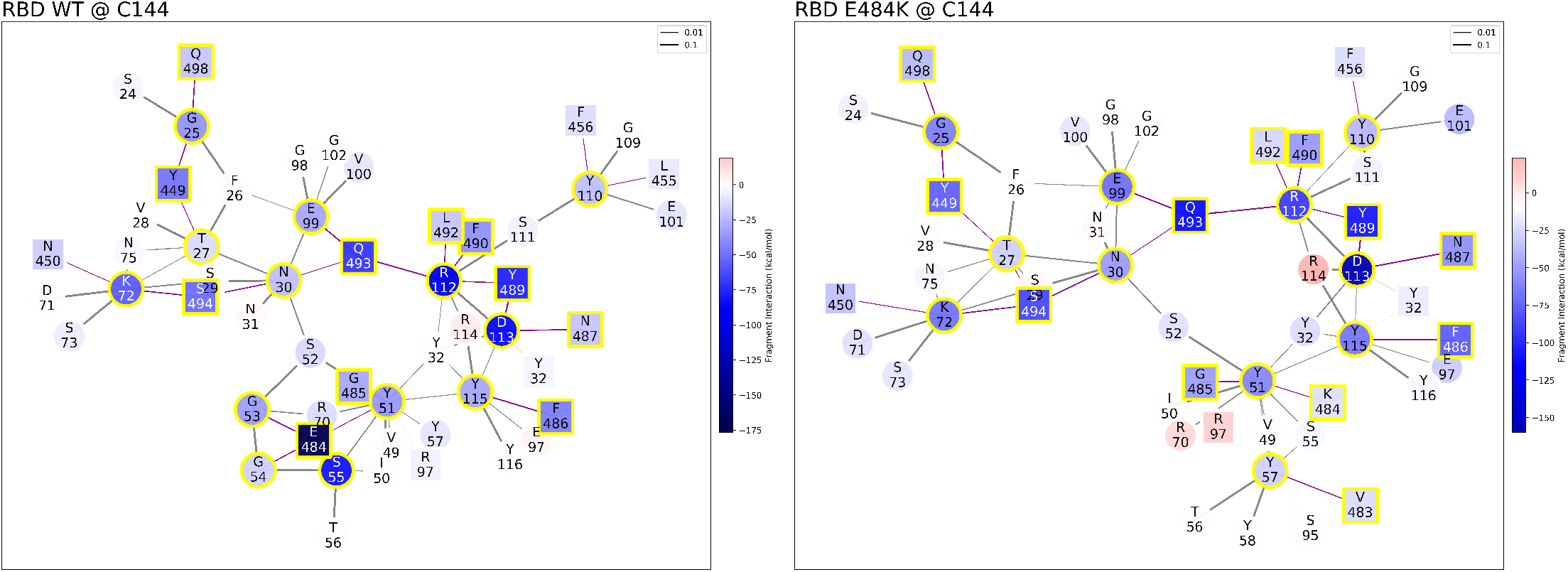
Mechanistic characterization of C144 binding to the Wuhan strain spike protein, and energetic changes as a result of the E484K spike mutation. Data are plotted on the spike primary structure (panel a) and on C144’s (panel b), considering the different bindings via the Wuhan spike (WT) and the mutated one (E484K). AA are represented by the letters, and numbered on the histogram’s horizontal axis. Histograms underneath the sequences represent the relative change in binding energy of the second row relatively to the first (Wuhan type strand). The bottom right histograms represent the overall binding energy of C144 with the Wuhan spike (right) and the mutated one (left) and its characterization as chemical or electrostatic. Interaction networks (RBD WT @ C144 to the left, RBD E484K @ C144 to the right) at the bottom represent the interface residues and their coordinated interactors: squares are spike residues and circles C144’s, their respective coloring is red for repulsive and blue for attractive energy, a yellow highlight represents interface residues. Bonds are purple when inter-molecular or black when intra-molecular. Interface residues are represented by red squares (repulsive) and blue squares (attractive) based on their effect to their counterpart, and highlighted by yellow squares when at the binding interface.

### A.2 *Ab initio* simulation shows how the E484K spike does not improve the binding energy to *Rhinolophus macrotis* ACE2 receptor (macACE2), compared to the Wuhan strain

**Fig A.2.**
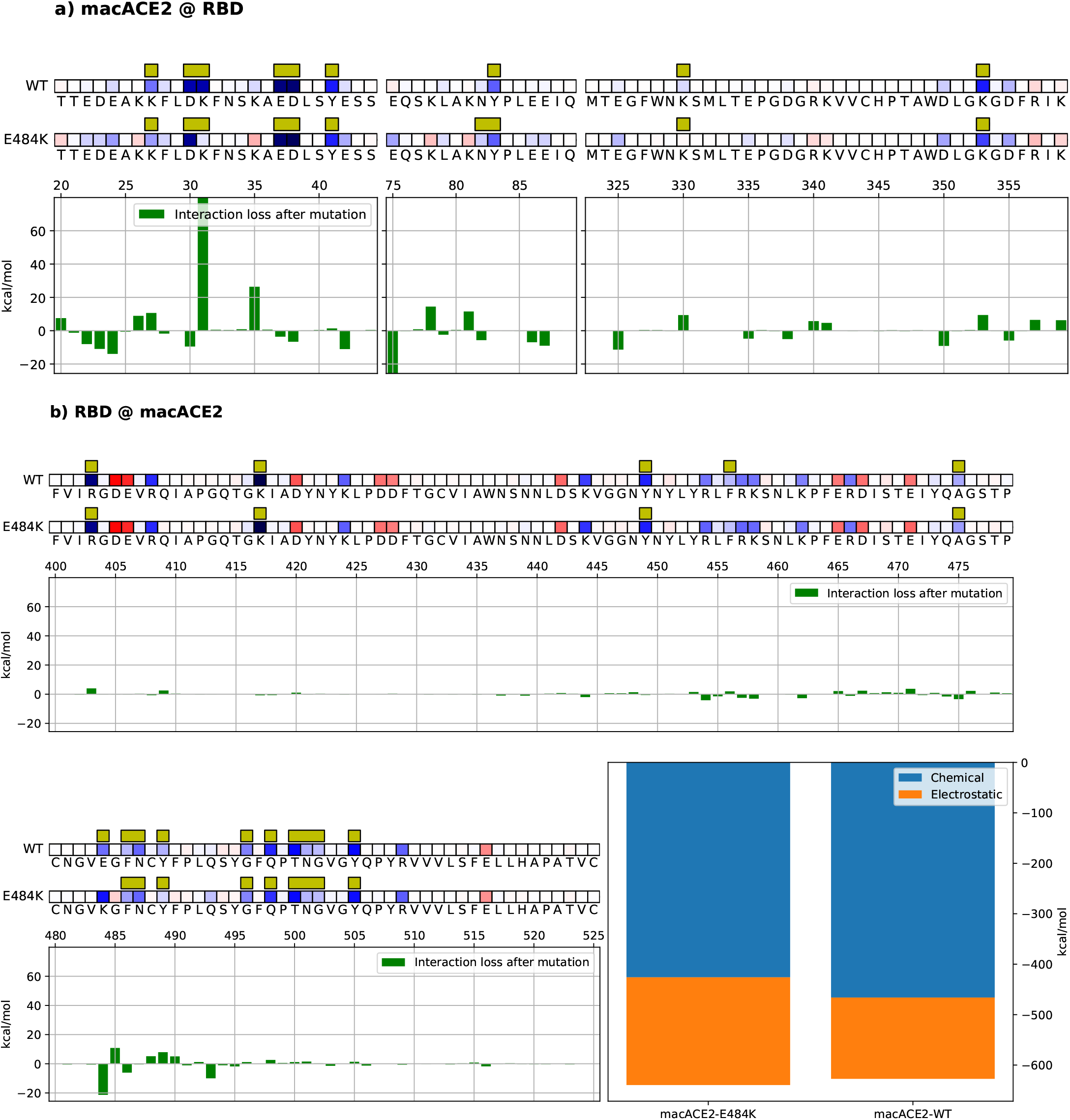
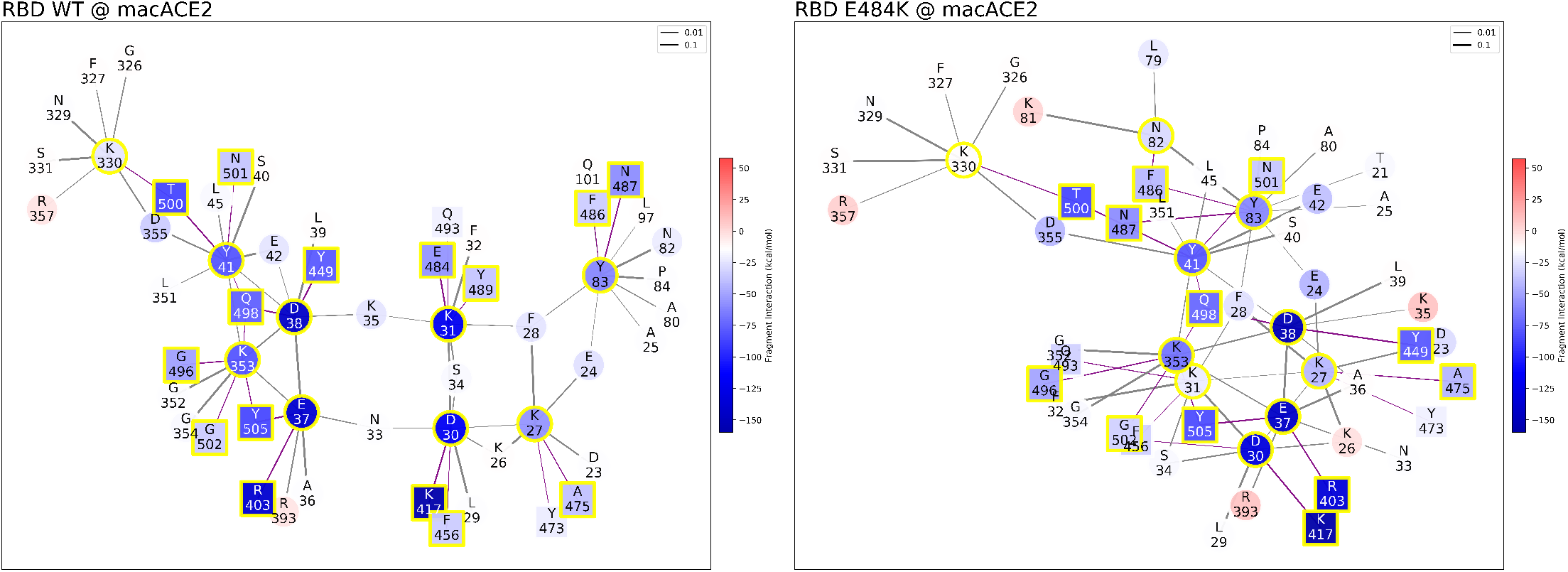
Mechanistic characterization of Wuhan and mutated (E484K) spike binding to macACE2. Data are plotted on the macACE2 primary structure (panel a) and on Spike RBD (panel b), considering the different bindings via the Wuhan spike (WT) and the mutated one (E484K). Amino-acids are represented by the letters, and numbered on the histogram’s horizontal axis. Histograms underneath the sequences represent the relative change in binding energy of the second row relative to the first one (Wuhan type strand). The bottom right histograms represent the overall binding energy of macACE2 with the Wuhan spike and the mutated one and its characterization as chemical or electrostatic. Interaction networks (RBD WT @ macACE2 to the left, RBD E484K @ macACE2 to the right) Bonds are purple when inter-molecular or black when intra-molecular, their width being related to the strength of the FBO between residues. Graph nodes are represented in red (repulsive) and blue (attractive) based on their effect to their counterpart, and highlighted by yellow squares when at the binding interface.

### A.3 Model comparison with preexisting experimental datasets on randomly generated libraries of hACE2 and viral spike mutants

We decompose the binding energy of the RBD with the ligand into per-residue contributions to describe the relative importance of each amino acid in the binding process. To see how these data compare with available experimental measurements, we generate a dataset of virtual point-mutations for the RBD-hACE2 system and compare the interaction energies with experimental affinity data obtained from existing high throughput random mutation experiments, which are available in the literature both for the Wuhan spike protein [32] and hACE2 [6]. Data are presented in Fig. A.3. As already pointed out in the main text, the two quantities are not directly comparable, as computational studies of protein-protein affinity require in-depth analysis of structural and thermodynamic contributions [48]. An example of discrepancy can be seen in the N501Y mutation, which appears strongly enhanced in the experiment and not in our model. This is possibly due to the fact that such mutation is associated to a structural rearrangement which would likely lead to different steric conformations. The E484K mutation stands out as an outlier in both the experimental dataset and our simulation, exhibiting an overall improved binding to hACE2. However, most mutations of the RBD residues lead to decreased affinity, indicating that the viral Wuhan spike is overall well-adapted to bind to hACE2.

### A.4 Details of the Fragmentation procedure

We here recall the main equations that define the energy decomposition schemes employed in the present work. We identify a system 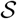 from the set of its atomic positions. A QM calculation of the system 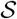 provides the density matrix 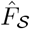. We associate to the atomic positions of the system 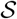 a set of ionic (electronic) charge densities 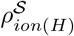, respectively. The expression of the total energy reads:

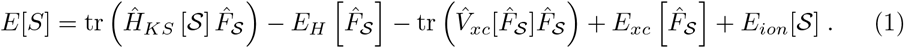

The DFT Hamiltonian is defined as 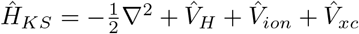 from the combination of the electrostatic potential provided by 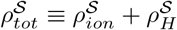, including the exchange and correlation term. Let us now suppose that our system is separated in two regions, which we call 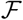 and 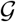, which in our example are associated to the RBD and the corresponding ligand.

When post-processing a QM DFT calculation, it is in theory possible to perform an analysis on an arbitrarily defined set of fragments. The challenge is to define those fragments in a chemically meaningful way, or, at least, to *quantify* the pertinence of a given fragmentation. To argument the choice of fragmentation scheme, in a previous study we introduced a measure of fragment quality called the purity indicator [23]. This measure is directly based on the density, being computed as the deviation from idempotency of the density matrix block associated with a given fragment. The interactions of a system may be described in the same framework by computing a quasi-observable called the “Fragment Bond Order” as a measure of the off-diagonal contributions of the density matrix. As described in [23], the Purity Indicator and Fragment Bond Order are computed directly from the density matrix. The Purity Indicator and Fragment Bond Order together measure the competition between a fragment’s internal and external interactions. This approach has recently been applied to generating graph views of proteins as well as to understand the interaction between proteins and solvent molecules [24].

We also assume that those two regions are associated to well-defined fragments with associated fragment projection operators 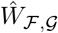. We can then identify, for instance 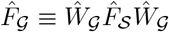. The operators 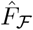 and 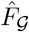 are then defined as the diagonal projection of the *full* density 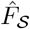 of the assembly into the regions associated to the subsystems 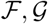. We define by 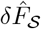 the off-diagonal term 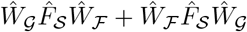

With these definitions, we can identify different terms contributing to the interaction of the fragments. Let us describe them separately.

- Electrostatic, Long-Range term: it is defined by the electrostatic interaction between the two fragments in their polarized state, ie. the ground state of the assembly.

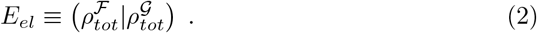 This is the only contribution which will have to be considered even when the 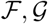 subsystems are spatially separated from each other. Such a term can be efficiently approximated by a sum of point-multipoles that are based on the description of the electron density. The electrostatic terms can be non-negligible even for fragments that are far apart, as the interaction which defines this term is long-ranged.
- Induction/Contact (Chemical), short-range term: it is defined by the trace of the block-off-diagonal part of the DFT Hamiltonian on the density of the total system. It reads:

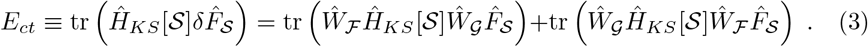 This term is a “contact” term, namely it is non-zero only for fragments which lie nearby, as the Hamiltonian and the density matrix are both represented by sparse matrices. Also, its interpretation is associated to the energy that the system would lose if both the fragments were considered as pure. As the purity indicator is associated to the fragment valence, this term is associated to the energy of the “chemical bond” between the fragments. For this reason, it is always associated to an attractive term, and its magnitude is directly correlated to the value of the fragment bond order. For compactness, dispersion vdW contributions coming from D3 terms [36] are included into such term, as the latter are also short-ranged. The Chemical term is always stabilizing, namely it only contains attractive interactions.

We have verified that the sum of these terms correlates very well with the QM interaction energy between the subsystems 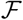 and 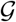, thereby proving that they capture the essential contributions of the interactions in our analysis [25].

Both these terms can be decomposed in per-fragment contributions; through the decomposition, we obtain an indication of the relative contribution that each fragment (in our case, each residue) provides to the interaction terms between the two assemblies. We represent those contribution in a network of interactions determined by the FBO quantity. (Absolute) purity values below 0.05 (see previous work [23, 24] for a justification of this threshold) indicate that the fragments are sufficiently pure and thus represent a good decomposition of the system. For the systems employed in this work, most of the amino acid lie within this range, with some exceptions (mainly Glycine residues) which reach a (yet acceptable) value of 0.07. For this reason, and to ease the discussion, we have preferred to stick with a usual fragmentation scheme based on aminoacids.

All the calculations are performed with the same DFT input parameters employed for the paper [25].

**Fig A.3.**
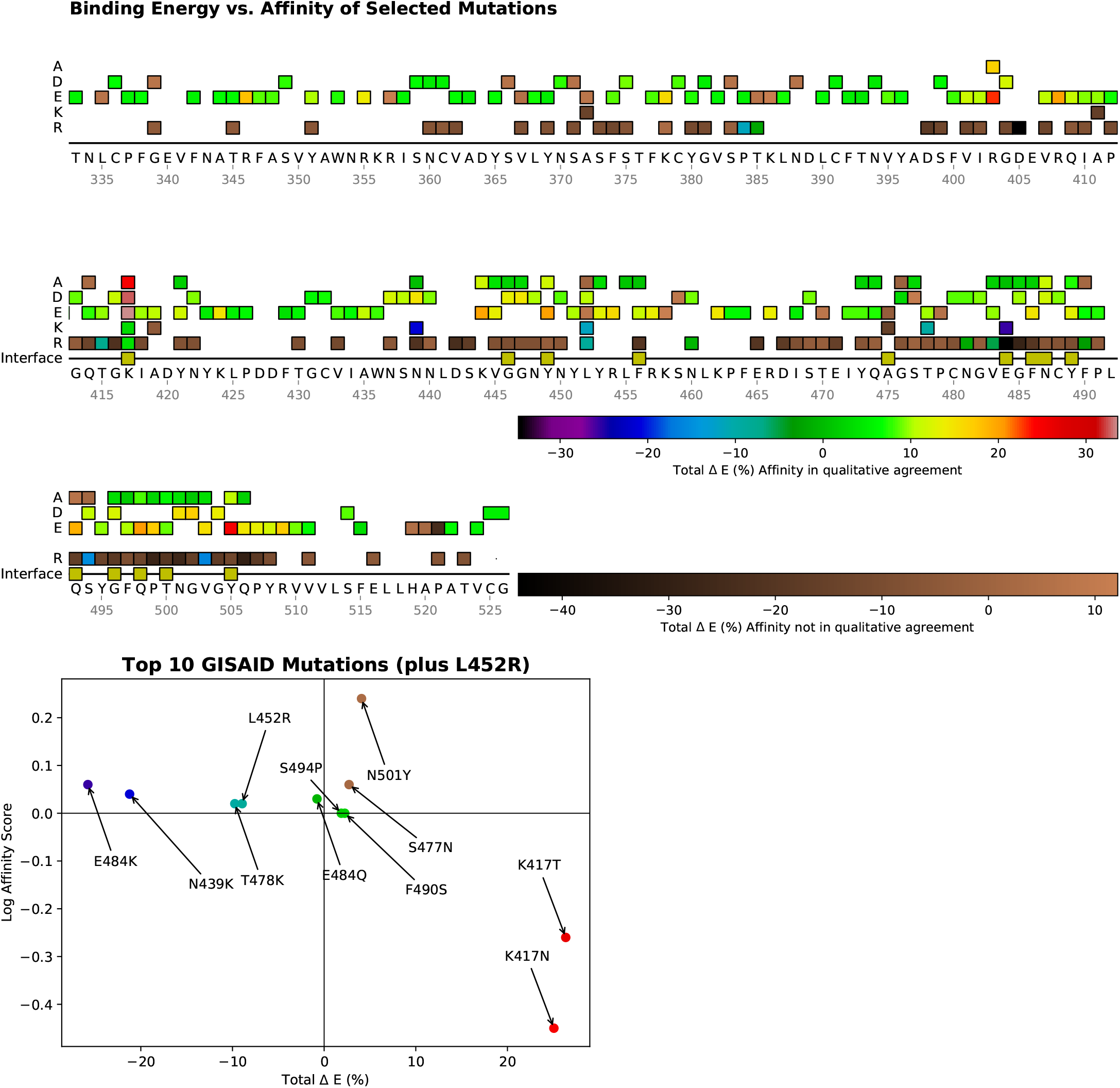
Comparison of *ab initio* simulations of virtual structures with available in vitro binding performance of mutant libraries. Spike mutations are represented for their affinity binding score towards hACE2 [32]. Negative score means depletion of the affinity of mutated sequence. Virtual crystals have been generated *in silico* and the binding strength simulated *ab initio*. The percent difference in binding strength is then calculated with respect to Wuhan strand, negative value indicating improvement of the binding. If both the *in vitro* and the *in silico* data represent a depletion or an enhancement, data are in qualitative agreement and colored with rainbow colors, from red to purple (errorbars and uncertainties have been neglected here), and with copper colors otherwise. The two different quantities are then represented in the plot (lower panel, horizontal axis for the *ab initio* binding, vertical axis for the affinity), by focusing on the top 10 GISAID [49] point-mutations (plus the L452R point-mutation) which are found in SARS-CoV-2 variants at the time of writing this contribution. The same color scheme is employed for the mutation represented in the sequence and for the plots. The two datasets have many points in accordance, more than 60% of the points being in qualitative agreement. Among all the tested mutations, E484K (purple point) emerges as the strongest *ab initio* binding performance in qualitative agreement with the experiment.

## Acknowledgments

We acknowledge useful discussions with Viviana Cristiglio, Michel Masella, Lorenzo Fontolan, Karla Ilic Djurjic. LG also acknowledges support from the MaX EU center of Excellence, and from French National computing resources (projects spe0011 and gen12049). BM and MZ were supported by an Ignite grant from Boston College and by an Award for Excellence in Biomedical Research from the Smith Family Foundation. This work used computational resources of the supercomputer Fugaku provided by RIKEN through the HPCI System Research Project (Project ID: hp200179).

## Author Contributions

Conceptualization: MZ, LG, MF, WJ, BM. Formal analysis: LG, WD. Funding acquisition: LG, TN, BM, WD. Investigation: MZ, LG, MF, WJ, BM. Methodology: MZ, LG, WD. Software: LG, WD. Supervision: MF, WJ, BM. Writing – original draft: MZ, LG, BM. Writing – review and editing: MZ, LG, MF, WD, BM.

